# Can the causal role of brain oscillations be studied through rhythmic brain stimulation?

**DOI:** 10.1101/2021.06.17.448493

**Authors:** Tanya Lobo, Matthew J Brookes, Markus Bauer

**Affiliations:** School of Psychology, University of Nottingham, University Park, NG7 2RD, UK; Sir Peter Mansfield Imaging Centre, University of Nottingham, University Park, NG7 2RD, UK

## Abstract

Many studies have investigated the causal relevance of brain-oscillations using rhythmic stimulation, either through direct-brain or sensory stimulation. Yet, how intrinsic rhythms interact with the externally generated rhythm is largely unknown. We either presented a flickered (60 Hz) visual grating or its correspondent unflickered stimulus in a psychophysical change-detection-task during simultaneous MEG-recordings to humans, to test the effect of visual entrainment on induced gamma-oscillations.

Notably, we generally observed the co-existence of the broadband induced gamma-rhythm with the entrained flicker-rhythm (reliably measured in each participant), with the peak frequency of the induced response remaining unaltered in approximately half of participants -relatively independently of their native frequency. However, flicker increased broadband induced-gamma-power, and this was stronger in participants with a native frequency closer to the flicker-frequency (‘resonance’), and led to strong phase-entrainment. Presence of flicker did not change behaviour itself, but profoundly altered brain-behaviour correlates across the sample: whilst broadband induced gamma-oscillations correlated with reaction-times for unflickered stimuli (as known previously), for the flicker, the amplitude of the entrained flicker-rhythm (but no more the induced oscillation) correlated with reaction-times. This, however, strongly depended on whether a participant’s peak frequency shifted to the entrained rhythm.

Our results suggests that rhythmic brain-stimulation leads to a coexistence of two partially independent oscillations with heterogeneous effects across participants on the ‘downstream relevance’ of these rhythms for behaviour. This may explain the inconsistency of findings related to external entrainment of brain-oscillations and poses further questions towards causal manipulations of brain-oscillations in general.

## Introduction

Numerous studies have shown that gamma-band oscillations, particularly in early visual cortex, are increased by bottom-up stimulation and modulated by task demands. Recent work in animals and humans suggests that gamma-band are particularly implicated in feed-forward processing hierarchical cortical circuits (M. Bauer, Stenner, Friston, & Dolan, 2014; Buffalo, Fries, Landman, Buschman, & Desimone, 2011; van Kerkoerle et al., 2014). Whereas initial suggestions that synchronization in the gamma-band may be a solution to the binding problem have received corroborative and contradicting evidence (Shadlen & Movshon, 1999; Singer, 1999), a particularly robust finding is that selective attention enhances gamma-band oscillations as a way to enhance the saliency of stimulus representations and the efficacy of neural signal propagation (M. Bauer, Oostenveld, Peeters, & Fries, 2006; Gruber, Muller, Keil, & Elbert, 1999; Womelsdorf et al., 2007). Indeed, several studies have found very robust correlative evidence that humans or animals perform better (e.g. faster reaction times) in trials where stimulus induced gamma-band oscillations are stronger, compared to when these are weaker (M. Bauer, Oostenveld, & Fries, 2009; Hoogenboom, Schoffelen, Oostenveld, & Fries, 2010; Womelsdorf, Fries, Mitra, & Desimone, 2006). Evidence for a mechanistic role of gamma-oscillations in the feedforwarding of information comes from a combined microstimulation and recording study showing that feedforward inputs ‘travel’ on a gamma-wave, whereas feedback inputs do so on alpha-/beta-waves (van Kerkoerle et al., 2014).

Furthermore, a good number of studies have attempted to assess the causal role of gamma-oscillations by artificially enhancing these through external entrainment, either through flickering stimuli at a gamma-frequency (F. Bauer, Cheadle, Parton, Muller, & Usher, 2009; M. Bauer, Akam, Joseph, Freeman, & Driver, 2012; Usher & Donnelly, 1998), through tACS (Kanai, Chaieb, Antal, Walsh, & Paulus, 2008) or through optogenetics (Sohal, Zhang, Yizhar, & Deisseroth, 2009), even though the latter allow for more specific control through cell-type specific activations. The rationale – and therefore underlying assumption -here is that artificially enhancing the amplitude of brain oscillations at the gamma-frequency range should enhance the mechanistic effects normally provided by the natural rhythm. However, it is not trivial to assume that this assumption is valid. Pikovsky, Rosenblum, & Kurths (2001) outline the different regimes of physical oscillators, contrasting, for instance, enslavement of oscillators by an external force with self-sustained oscillators (the so-called limit-cycle). Indeed, the emergence and strength of gamma-oscillations in the brain appears to depend on a finely tuned balance between inhibitory and excitatory neurons and generally involves the coupling of many neurons (Tiesinga, Fellous, Salinas, Jose, & Sejnowski, 2004). These interactions are fundamentally different from pulsed excitation of a very broad range of excitatory neurons by an external force (e.g. bottom-up stimulation by a physical stimulus, as in the work presented here and numerous other studies). Moreover, it is well known that the amplitude of the entrained brain rhythm depends crucially on the applied frequency, with the typical 1/f dropoff. In other words, higher frequencies, such as the gamma-band, often classified as approximately 30-120 Hz and in human calcarine sulcus most typically found at frequencies of about 60 Hz (Hoogenboom, Schoffelen, Oostenveld, Parkes, & Fries, 2006; van Pelt, Boomsma, & Fries, 2012), show considerably smaller amplitudes in response to flickering stimuli than lower frequencies below 30 Hz (Herrmann, 2001). One study investigating this in Macaque monkeys (and humans) has revealed that V1 neurons are capable of following rhythms at frequencies up to 100 Hz, but only a subset of these are actually entrained (Williams, Mechler, Gordon, Shapley, & Hawken, 2004), leaving the question of the impact entrainment at such frequencies might have on visual processing throughout the brain. For tACS, the question has been raised to what extent this type of brain stimulation is effective at all in modulating neural activity (Liu et al., 2018), although there is hard evidence that clearly shows the effectiveness of tACS to modulate neuronal activity (Johnson et al., 2020) and many more studies providing clear evidence in favour of it. Nevertheless, the question remains how the externally applied rhythm truly interacts with those neurons usually involved with generating intrinsic gamma-rhythms.

Several studies have used flicker-paradigms to probe the hypothesis that gamma-oscillations support neurocognitive processes such as feature binding or attention. Usher & Donnelly (1998) found that gamma-flickered stimuli at 50 Hz can facilitate binding of stimuli into a coherent object, by inducing stimulus dependent synchrony. However, others have provided conflicting evidence (Kiper, Gegenfurtner, & Movshon, 1996). F. Bauer et al. (2009) claimed to have shown that flickering stimuli at 50 Hz enhanced stimulus saliency by attracting attention at a subliminal level. They were flickering a stimulus in one particular spatial location embedded amongst two distractor stimuli and showed an increase in stimulus-change detection. However, in a subsequent study, van Diepen, Born, Souto, Gauch, & Kerzel (2010) have found conflicting results, suggesting that the positive findings of (F. Bauer et al., 2009) study may have been due to potential confounds. M. Bauer, Akam, et al. (2012) have probed the effect of flicker stimuli (60 Hz) on visual signal propagation and selective attention / binding processes, found no effect of flicker stimuli on stimulus saliency or stimulus selection. Taken together, these studies show that the use of high-frequency flicker provide at the very least ambivalent results and somehow question the efficacy of entraining gamma-oscillations, at least by flicker stimuli.

Here, we set out to investigate the interaction of an externally entrained rhythm by flickering visual stimulation with intrinsic gamma oscillations. We used a paradigm similar to some of the cited studies that have shown that gamma-oscillations are implicated in selective attention processes and visual signal propagation. Specifically, we used a stimulus change detection paradigm where a high-contrast sinusoidal grating was presented for an extended period (4 s) and participants had to detect a change in the spatial frequency that occurred unpredictably, following a uniform probability distribution. The grating was either static or it was flickered at 60 Hz, with the contrast level in the two conditions being perceptually matched in a previous session and otherwise being of identical physical properties.

We addressed the questions of whether the 60 Hz flickering stimulus would

1. enhance gamma-oscillations and, more specifically, whether the flicker stimulus would entrain brain activity at its frequency whilst suppressing broadband-induced oscillations (as they are usually observed with such stimuli) or whether the two rhythms might co-exist,
2. facilitate behaviour in terms of detection performance and reaction time when compared to the statically presented grating. Additionally, we asked,
3. whether the effect of the flicker depended on the intrinsic gamma-peak-frequency of a particular individual and specifically its distance from the external 60 Hz rhythm (resonance)
4. if flicker and intrinsic broadband rhythm were to co-exist, which rhythm appears to be more instrumental for the behaviour of participants by way of correlative relationships.

## Methods

Fourteen healthy participants, naïve to the purpose of the experiment, were recruited for the experiment from a database at the School of Psychology at the University of Nottingham in agreement with the local ethics committee for taught projects (as part of a BSc degree in Psychology and Cognitive Neuroscience). All fourteen were paid for their participation in the study (£15 for one psychophysical training session and one MEG session). Participants were excluded on the basis of a close/personal family history of epilepsy due to the flickering nature of the stimuli. They were also asked to ensure that they had no irremovable metal in/on the body. Since the LCD monitor in the MEG scanner would be at a fairly large distance from the participant, potentially interested participants with poor visual acuity were excluded from the study.

The experiment was run using Presentation (Neurobehavioral Systems) software. The stimuli presented were black and white sinusoidal gratings at a spatial frequency of approximately 2.28 cycles per degree of visual angle, with a size of approximately 6 degrees per visual angle on a black background (Figure 1). Stimuli were presented on a VPixx® LCD monitor, with a refresh rate of 120Hz, either constantly, on each individual frame (static condition), or in alternation with a black screen (60 Hz flicker condition). The same number of trials of flicker stimuli and non-flicker stimuli were presented within a block of 18 trials (288 trials in total). The screen was located at a distance of 2.13m, in both the psychophysical and the MEG session, in a darkened room (during MEG that was the magnetically shielded room, MSR) and was adjusted to have a linear contrast gain.

**Figure 1.**
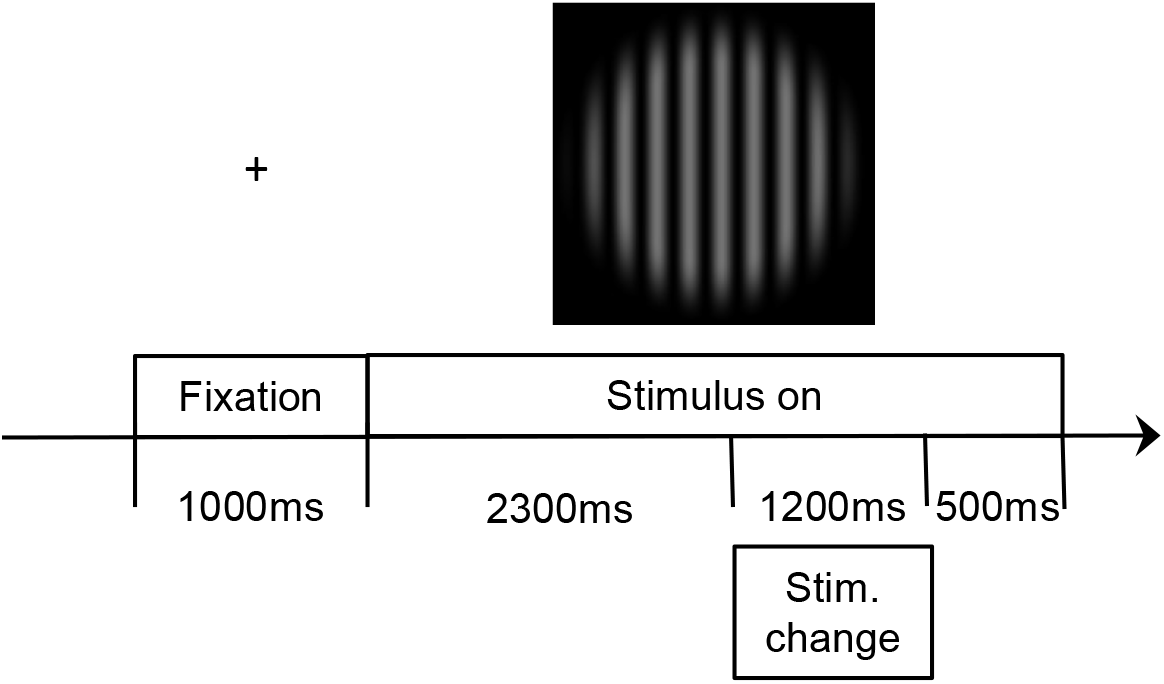
Main experimental task. A fixation cross was presented for 1s, followed by the onset of either a central, flickering (60 Hz) full contrast grating or a contrast matched static grating appeared for a total period of 4s (fixed). On 80% of trials a spatial frequency change occurred between 2.3 and 3.5s following stimulus onset, following a uniform distribution. Participants had to press a button as fast and accurately as possible.

**Figure 2.**
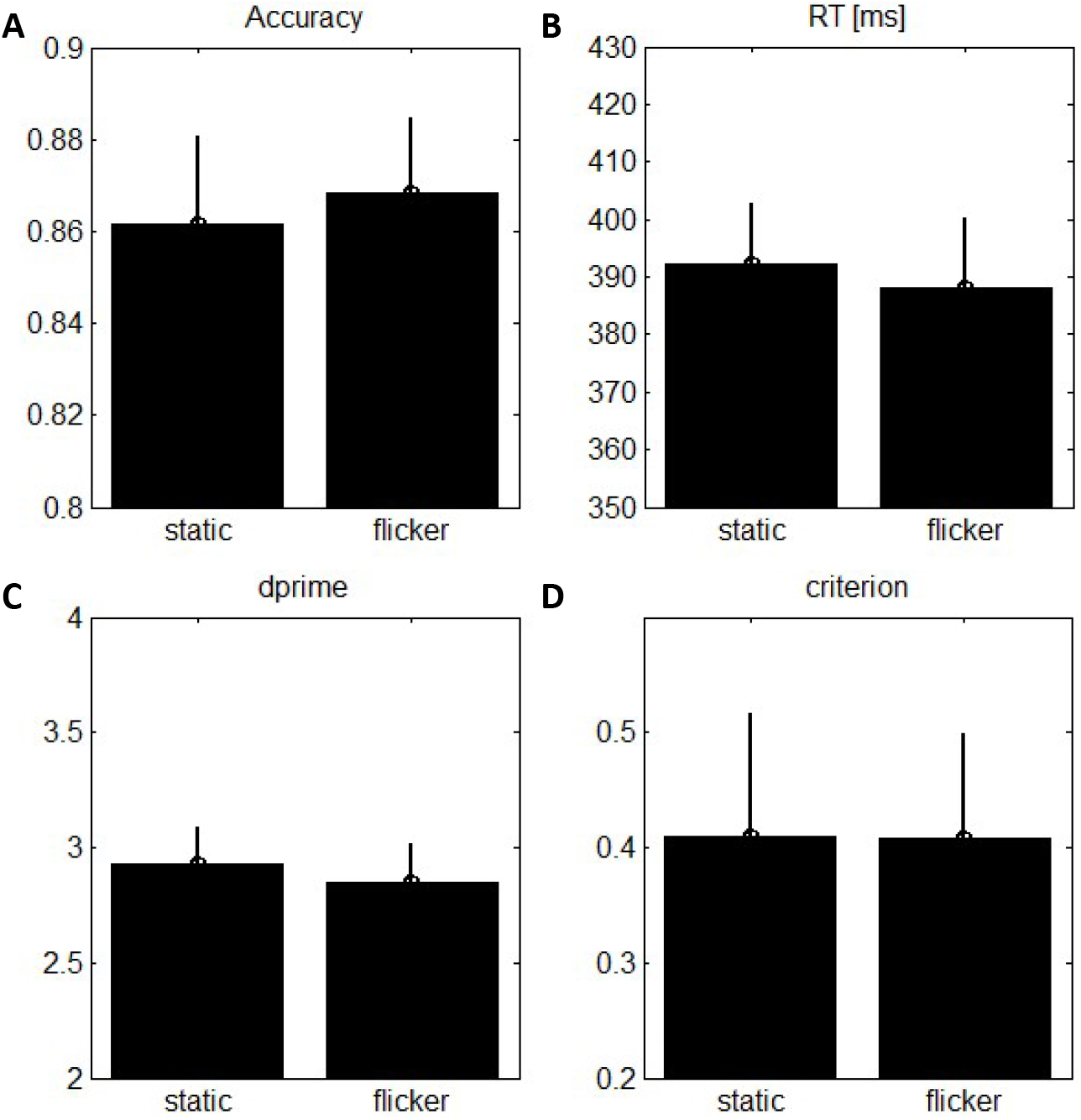
Behavioural results for the detection task. **A)** Proportion of correct responses (hits and correct rejections). **B)** Reaction time after spatial frequency change. **C)** Signal-detection theory (SDT) d’ parameter. **D)** Response criterion (SDT). No differences between conditions approached statistical significance (all paired t-tests: −0.8 < t < 1.1; p > 0.2, one-sided). All errorbars denote standard error of the mean (SEM).

### Procedure

After having given informed consent and having confirmed that neither them nor close family members were suffering from epilepsy (nor the participants having neurological disorders or being under medication), participants underwent a training session lasting thirty minutes, prior to the experimental session in the CTF Omega 275 channel MEG scanner (that took place on a different day). The training session was necessary to familiarize participants with the task and to adjust, for each participant individually, the contrast level of the static grating stimulus to the 60 Hz flicker grating.

Since the flickering stimuli were presented on every alternating frame, their effective contrast/luminance is reduced and in order to account for these contrast differences, prior to the start of the behavioural training session, each participant was presented with both static and flicker stimuli and asked to subjectively match the luminance/contrast of the static grating to the flickered grating using the ‘up’ and ‘down’ arrow keys. This contrast level matching was done 9 times, after which a mean value for the static stimulus was calculated for each participant and subsequently used for the training session and main experiment. Such a subjective criterion was used given that different neurons are likely to have different transfer functions with respect to temporal properties and nonlinearities in contrast gain.

To adjust the spatial frequency change, an adaptive staircase procedure was used, such that after blocks with less than 80% accuracy, the spatial frequency change was increased, whereas after blocks with more than 90% accuracy, the spatial frequency change was decreased.

For the main experiment, as well as the training task, at the beginning of each trial a fixation dot was presented for a period of 1 second. The grating stimuli were then presented for a period of 4 seconds. The spatial frequency change could occur unpredictably between 2.3 and 3.5 seconds in order to ensure sustained attention from participants but only occurred in 80% of all trials. Participants reported the spatial frequency change with their right index finger.

For the MEG session, participants were comfortably seated in the MSR and head localisation coils were then placed at the nasion and pre-auricular points of the skull for head localisation. The main task was divided into 16 blocks consisting of 18 trials each. In between each trial, a fixation dot was presented at the centre of the screen. After each block, a black screen was presented during which participants could rest their eyes and continue the task when ready via button press. A photo-diode was placed into one of the corners of the screen and its signal fed into the ADC (analog-digital-conversion) channels of the MEG, thus in synch with the MEG sensors. A square was shown at the position of the photio-diode (hidden away from the participant) in synchrony with the flickered grating, allowing to check for irregular or missed frames and serving as a reference signal for flicker-entrained brain responses. MEG was measured at a sampling rate of 600 Hz, with a hardware anti-aliasing filter at ¼ of the sampling rate (150 Hz).

### Processing of MEG data

Artefact removal was conducted using SPM8 (Litvak et al., 2011) and FieldTrip (Oostenveld, Fries, Maris, & Schoffelen, 2011). A major source of artefacts in the MEG scanner are eye movements. Data was epoched 200-300ms around the eye blink in the main data and a principal component analysis was run in order to extract components which maximize variance out of the data. A spatial pattern of eye blinks was identified and regressed from the data using signal space projection; remaining artefacts were removed according to a summary statistic showing the total variance in individual trials.

To maximise signal to noise ratio (SNR), all further analyses were conducted in source space, using a beamformer transform. Previously, we had run the same analyses in planar gradiometer sensor space, and while these analyses returned qualitatively the same effects with respect to the main effects, the correlations with behaviour were not statistically significant there. To this end, we used SPM8 for constructing a cortical sheet representation of the brain (M. Bauer, Kluge, et al., 2012; M. Bauer et al., 2014) based on the canonical MNI brain as implemented in SPM8 (Litvak et al., 2011). Beamformer coefficients (Van Veen, van Drongelen, Yuchtman, & Suzuki, 1997) were computed on the basis of the covariance-matrix of high bandpass-filtered data (40 – 150 Hz, please note that a bandpass-filter is not rectangular and will also pass substantial signal e.g. at 30 Hz) as well as the leadfield (taking individually measured head-positions in the sensory array into account). For frequency analysis, the pre-processed raw (sensor level) data were projected through beamformer coefficients corresponding to 413 individual cortical grid points (to make the data more treatable, (M. Bauer, Kluge, et al., 2012; M. Bauer et al., 2014) and all subsequent analyses were carried out on these local estimates of cortical brain activity.

### Frequency analyses

Two separate frequency analyses were calculated on these time series: 1) for the induced power (30-130 Hz) broadband gamma-band-response, a multi-taper analysis (M. Bauer et al., 2006; Mitra & Pesaran, 1999) was used with 0.4s long windows in steps of 0.1s and for frequencies in steps of 2.5Hz with 20Hz smoothing (resulting in 7 tapers); 2) for the steady state narrow band (60 Hz) response, the entire 4s period was Fourier-transformed, using a Tukey window for optimal spectral resolution (0.25 Hz).

These two separate analyses were conducted for the following reasons: Firstly, steady-state-induced responses by flickering stimuli tend to be very narrowband, typically with a bandwith of considerably less than 1 Hz (Kamphuisen, Bauer, & van Ee, 2008; Srinivasan, Russell, Edelman, & Tononi, 1999), benefitting considerably from a high spectral resolution (i.e. low smoothing) for obtaining a good SNR. Secondly, visually induced gamma-oscillations show the very opposite characteristic: namely of being very broadband in nature (M. Bauer et al., 2006; Fries, Reynolds, Rorie, & Desimone, 2001; Hoogenboom, Schoffelen, Oostenveld, Parkes, & Fries, 2005; Muthukumaraswamy & Singh, 2013; Siegel & König, 2003) and therefore benefitting considerably from greater smoothing (i.e. lower frequency resolution) in the frequency domain (Mitra & Pesaran, 1999).

We initially looked at the absolute power spectra for each of these analyses, but subsequently also investigated

1. the phaselocked response to the flicker-stimulus in the narrowband
2. the phaselocking to the flicker-stimulus in the broadband induced gamma response
3. the broadband induced response after removing the narrowband 60 Hz (flicker) response to assess the wider consequences of the flickered stimulus on brain activity after removing its directly entrained activity.

For 1) the same data were used as for the power analysis, whereas for 2), we computed the complex Fourier coefficients of 1s long windows, to optimise SNR at the expense of temporal resolution with 20 Hz smoothing (resulting in 19 tapers for each of which the complex Fourier coefficients were calculated separately). For the narrowband data we calculated both the a) absolute value of the averaged complex Fourier-coefficients, to obtain the magnitude of the phase-locked (or evoked) response and b) the phaselocking-value (Lachaux, Rodriguez, Martinerie, & Varela, 1999) which was obtained as the magnitude of the averaged, complex, normalised Fourier-coefficients (where the complex Fourier spectra of both time-series were each normalised on their magnitude on an individual trial basis), such that the magnitude of the individual trial Fourier estimates was always one and they only differed in their relative phase. This metric purely reflects the degree of phase-entrainment by the flicker-stimulus, in absence of signal amplitude. For the broadband data we were only interested in the latter, the phaselocking-value.

For the removal of the direct consequences of the 60 Hz flicker, both for simplicity and to be on the more conservative side, we simply used a bandstop-filter from 59.75 to 60.25 Hz to remove the total power (i.e. not only the phaselocked response was subtracted but the whole narrowband 60 Hz signal) from this direct flicker response. The rationale for this is that, if subtracting a signal of average phase (the phaselocked or evoked response) from a signal of random phase, this will not just effectively remove a phase-locked signal but would also introduce new artefacts into the data.

Whilst the VPixx® monitor had very good synchronization properties, there were occasional frame-lapses (as identified from the photo-diode signal) in on average 2.1% of all trials (range across subjects: 0-7.3%, where the highest number is presumably due to a temporarily unplugged photodiode at the beginning) and these were excluded from all analysis where stimulus phase mattered (in particular, the evoked / phase-locked flicker oscillations analyses, as well as phase-locking analyses on both broadband and entrained flicker rhythm (Figures 3, 4 and 5).

**Figure 3.**
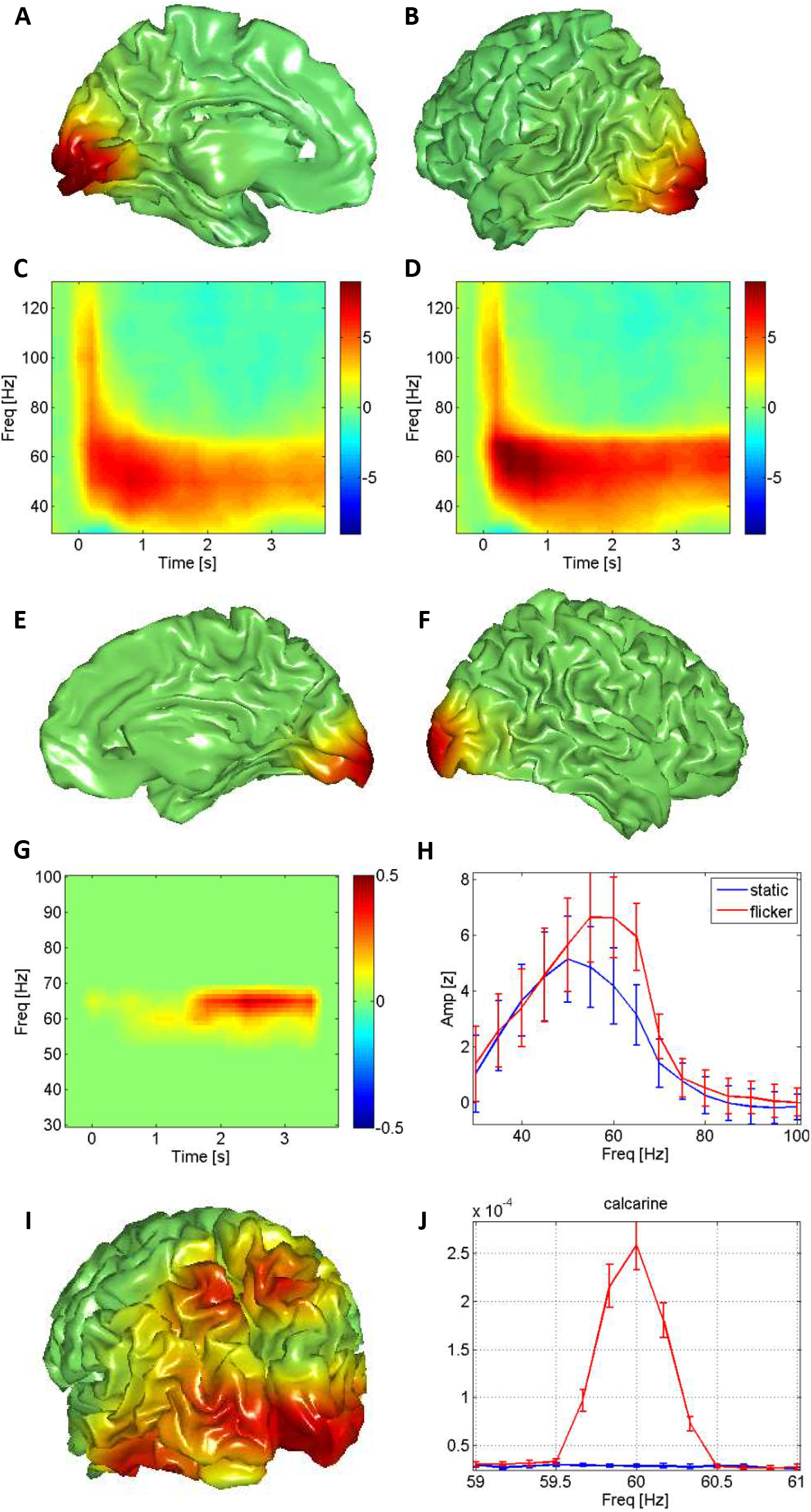
Effect of flicker on MEG activity. **A), B)** Topography of induced gamma-band effect (static condition only) from medial and lateral perspectives, respectively. Shown is the statistical comparison of post-stimulus power against baseline, thresholded for multiple comparisons (p<0.05). The colourscale represents arbitrary units (see text) **C)** Time-frequency-representation (TFR) of the changes against baseline (t-statistic) in significant cortical grid points (A), in static conditions. The colourbar represents t-values. **D)** Same for 60Hz-flicker. **E), F)** Thresholded topography of statistical effect of 60 Hz flickered stimuli vs static condition, medial and lateral view, respectively (colourbar arbitrary units, see text). **G)** TFR of the thresholded statistical comparison ‘flicker’-’static’. Colourbar represents arbitrary units, see text. **H)** Spectral profile in significantly modulated grid points for flicker and static. **I)** Topography of the thresholded statistical comparison between flicker and static on phaselocked narrow-band responses. **J)** Spectral profile of flicker (red) and static (blue) activity in significantly modulated grid-points shown in (I). All errorbars represent SEM.

**Figure 4.**
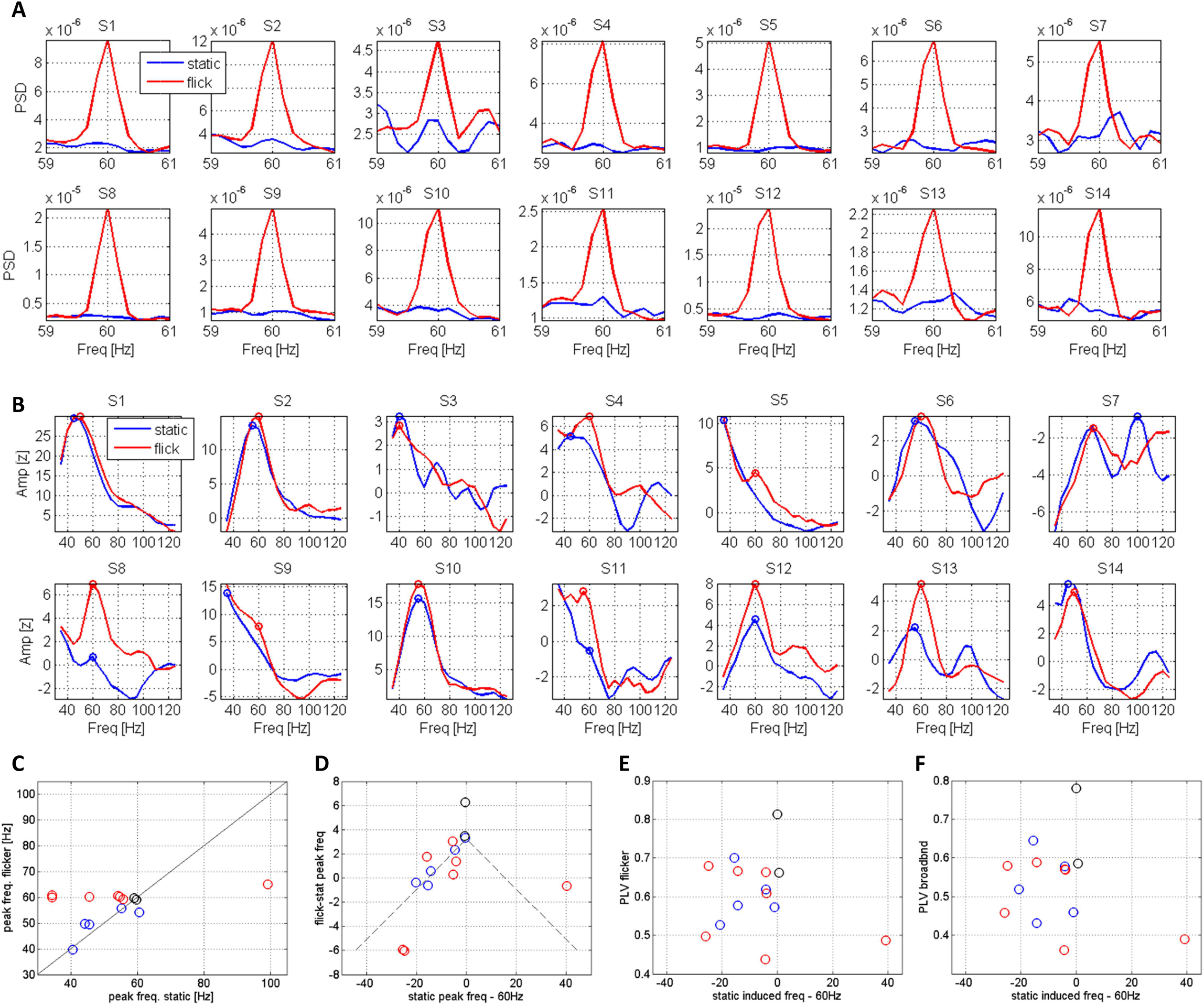
Individual spectra and patterns of ‘entrainment’. **A)** Power-spectra for narrowband 60 Hz responses in occipital cortex for flickered and static stimuli. Shown is the absolute magnitude of the power-spectrum, not only the phaselocked component. **B)** Spectra of broadband induced responses in occipital cortex for flickered and static stimuli, for each participant, corresponding numbers to A). Circles mark the estimated peak-frequencies of the spectra. **C)** Peak frequencies of broadband induced responses in flickered and static conditions (as shown in B). These reveal two distinct patterns: one (marked in red) that show a shift from the static peak frequency (not 60 Hz) towards the flickered frequency (60 Hz); one (marked in blue) where the peak frequency either does not change by flicker or moves (slightly) away from 60Hz. Black circles represent those participants whose peak-frequency in static condition was 60 Hz and remained unchanged. The colour scheme in figures D-F follows this pattern, see also Fig. 5F. **D)** Peak amplitude difference of broadband induced responses between flickered and static conditions as a function of peak-frequency (expressed as a difference from 60 Hz), showing resonance pattern; the black line represents the fitted linear regression. **E)** Phaselocking-value of narrowband 60 Hz-responses as a function of static peak-frequency. **F)** Phaselocking-value of broadband gamma-responses (taken at 60 Hz) as a function of static peak-frequency. No dependency on either static peak frequency or whether the peak frequency in flickered state shifts are evident.

**Figure 5:**
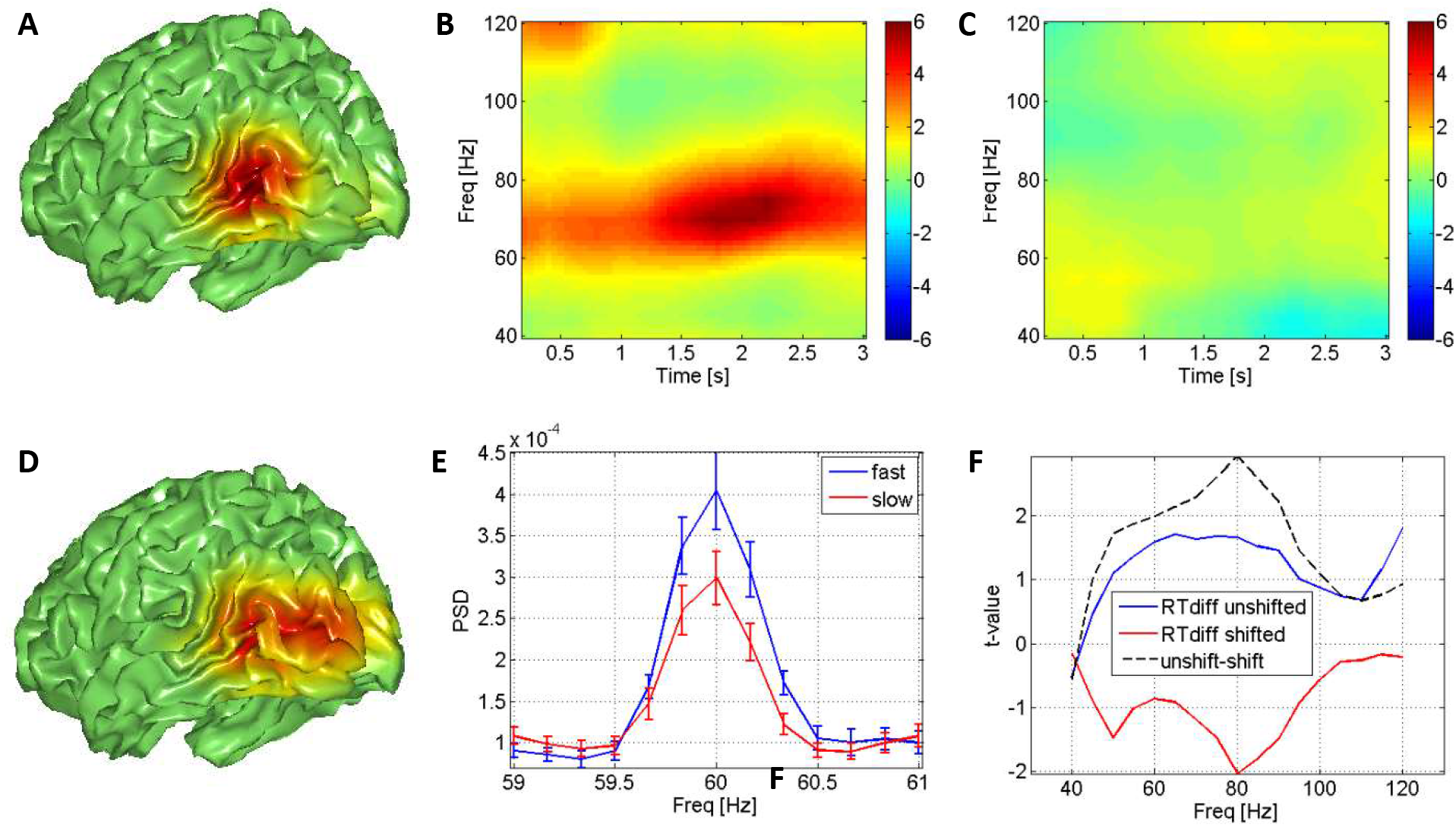
Behavioural correlates of induced and entrained responses. **A)** Topography of thresholded statistical comparison between *fast and slow trials in static condition* on broadband induced gamma-activity. Colourscale represents arbitrary values. **B)** TFR of same comparison in significantly modulated cortical grid-points. Colourbar shows t-values. **C)** TFR of the same comparison (fast-slow) in 60Hz flickered condition (across all participants): no effect visible. **D)** Topography of thresholded statistical comparison between fast and slow trials (like A) in flickered condition on narrowband phaselocked activity. **E)** Spectra of activity in fast vs slow responses in cortical grid points as shown in (D). **F)** shows the comparison of (C), i.e. fast vs slow trials in static condition on broadband induced gamma responses in the flicker condition, for two subpopulations collapsed across the time dimension: in red those participants that shifted their (broadband) gamma peak frequency to the flicker rhythm (see Figure 4 C) and in blue those that did not shift. Those participants, whose ‘intrinsic gamma rhythm’ did not shift by the flicker stimulus fast trials, had enhanced gamma-power compared to slow trials (as in Fig 5B but not 5C) whereas those participants, whose gamma-peak frequency got entrained by the flicker, this effect did not prevail. The black line shows the direct t-statistic between those groups.

The analyses on phase-locked responses for the flicker stimulus thus only includes trials where the entire 4s of stimulation period shows regular periodic flicker.

To estimate the individual peak frequency responses, a semi-automatic procedure was used.

An ROI was defined in the calcarine sulcus, where the spatial peak of the induced gamma-response was clearly located. The spectra of baseline-corrected (absolute) power modulations (t-values of each peri-stimulus window for each frequency against a baseline window from 400 to 0ms, i.e. just one temporal window, prior to stimulus onset) was therefore averaged from 400ms post-stimulus (to cut out the very broadband, high frequency early evoked response and focus on the quasi-stationary part thereafter, see Figure 2B, C) until the end, 3.6-4s post-stim.) and the two spectra, for flickered and unflickered stimulus were displayed on top of each other. The maximum of each power spectrum was chosen as default, but the location of this peak could be manually altered by the researcher (M.B.), using the following rule: by default, the frequency of the maximum of the entire spectrum was chosen; if, due to the 1/f signal dropoff, the maximum was the 35 Hz power estimate, the next largest peak was chosen. If no such maximum was present (i.e. the spectrum fell monotonously), the 35 Hz estimate was chosen. The individual peak-locations, together with their spectra, are displayed in Figure 4A (circles reflect the estimated peak-frequencies that entered the analysis). We chose against a fully automatic function or fitting mathematical functions to the spectra due to the (well-known) heterogeneity of spectral shapes that can, in our opinion, not be grasped well by any particular mathematical function, at least not without reasonably constraining the parameter-space which would otherwise likely lead to an overfit.

### Statistical analyses

Cluster-permutation tests (Maris & Oostenveld, 2007), simultaneously correcting for multiple comparisons across space (cortical grid points in 3-dimensional space), time and frequency were run for all of the following comparisons: 1) stimulation against baseline on broadband power-spectra for unflickered gratings against baseline (Fig 3A-D); 2) the same for flickered grating against baseline on induced broadband responses (giving virtually identically shaped clusters, not shown); 3) flickered grating vs static grating on broadband induced responses (Fig 4E-G); 4) flickered gratings vs unflickered gratings on phase-locked narrowband responses (these had no time dimension, given the 4 second window and thus no correction for multiple comparisons over time), Fig 3I; 5) fast vs slow trials in unflickered trials on broadband induced responses (Fig 5A); 6) fast vs slow trials in flickered trials on broadband induced responses (no significant clusters observed, Fig 5C); 7) fast vs slow trials in flickered trials on narrowband phase-locked responses (Fig 5D). These tests were conducted more or less in analogy to previously published results (M. Bauer et al., 2014). Here, however, they were calculated for the whole brain (rather than combined hemispheres), with the following parameters: Neighbouring cortical grid points had to be within 25mm distance, two-sides tests were run at univariate significance (or clustering-threshold) of p < 0.01, and a minimum of 3 neighbouring grid points were required. For all analyses except 4), to test for the phaselocked responses in the flicker vs static responses, 1000 permutations were computed. For the latter this was expanded to 4000 repetitions since the significance value of the cluster was so low. The omnibus significance criterion was, however, set for all tests at p < 0.05 two-sided.

To separate fast from slow trials, the fastest 16.7% of trials (in each condition) were contrasted with the slowest 16.7% of trials, see (M. Bauer et al., 2014; Hoogenboom et al., 2010; Womelsdorf, Fries, Mitra, & Desimone, 2005).

Finally, to create the brain topographies in Figures 3 and 5 (presented on ordinary MNI brain surfaces, rather than a heavily downsampled version of it), the statistical results for the 413 cortical gridpoints were interpolated to a standard cortical mesh consisting of 8196 grid points (as standard in SPM), by convolving the downsampled grid with a three-dimensional Gaussian Kernel with a standard deviation of 1cm and total extent of 3.5 standard deviations (i.e. 3.5 cm). This obviously leads to some blurring of the results but it is easier for novices to appreciate the topography this way. Importantly, no statistical analyses were conducted on such inflated data, this is just for graphical display purposes.

## Results

### Behavioural Results

Behaviourally, there were no significant differences found between statically presented and flickered gratings, neither in the accuracy (hit-rate) of detecting the stimulus change (t=-0.41, p=0.69) nor in reaction times (t=1.07, p=0.30), nor in signal detection parameters (d’ and criterion, all p > 0.2). The results are graphically shown in Figure 2.

### Electrophysiological Results

#### Induced gamma-band responses

Given the different spectral properties of steady-state-evoked responses to flickering stimuli and visually induced gamma-responses to grating-stimuli, we analysed the data with two separate frequency analyses capturing the narrow-band and broad-band aspects of these (see methods section). Figure 3 shows the stimulus induced and evoked responses. The topographies in Figure 3A and B (medial and lateral views of the left hemisphere for illustration) show the topography of the statistical comparison (power enhancement) of the post-stimulus-period to baseline (averaged over significantly modulated frequencies and time points) in the non-flickered condition (p = 0.025). The topography for the flickered grating (p = 0.02) looks virtually identical and is therefore not explicitly shown (see Fig 3E and F for the direct comparison though); both effects are bilateral with no obvious hemispheric asymmetry (not tested) and clearly localise, despite the lack of individual anatomical MRIs, into the calcarine sulcus. This result was thresholded for multiple comparisons across space, time and frequency. Figure 3C shows the time-resolved spectrum of this statistical comparison (in those grid points that are significantly modulated, see Fig 3A and B) for the unflickered condition, and Fig 3D shows it for the flickered condition. Note that there is an initial very broadband high-frequency enhancement of activity in frequencies of up to approximately 120 Hz which then transforms into a more ‘stationary’ response with a lower and more narrowband (still very broad, though note that these spectra have consideral spectral smoothing) oscillatory response. This broadband oscillatory response in the flicker condition shows a substantially enhanced amplitude compared to the non-flickered condition.

#### Enhancement of induced gamma-band oscillations by the flicker stimulus

The direct comparison between the responses in the two conditions (‘flicker’ -‘non-flicker’) is shown in Figures 3E-H. Figures 3E and F show the thresholded topography of this statistical comparison (p = 0.009, corrected), in analogy to Fig 3A-B, now for the right hemisphere, medial and lateral view, respectively. This topography shows a focal maximum of the induced gamma-enhancement by the flicker-stimulus at the posterior pole of occipital cortex, near the calcarine sulcus. Figure 3G shows the (thresholded) time resolved spectrum of the statistical comparison on broadband power in the statistically significantly modulated cortical grid-points (Fig 3E-F). This shows that there is a statistically significant enhancement over a wider frequency range from about 55-70 Hz, where the statistically most significant modulation occurs at approximately 65 Hz (note that this does not necessarily coincide with the biggest amplitude difference). Figure 3H shows the baseline-corrected amplitude spectra of flickered and static conditions averaged across time in the same cortical grid-points, confirming that the majority of power-enhancement is above 60 Hz (this is not an artefact).

We replicated this analysis with the removal of the neural activity directly caused by the flicker-stimulus. Rather than subtracting only the neural activity that is phaselocked to the flicker-stimulus, we used a bandstop-filter from 59.75 to 60.25 Hz and repeated the above analyses. This replicated the findings for the broadband stimulus induced activity, for both static and flickered conditions rather well (p = 0.034 for both static and flickered condition, separately, see Suppl. Fig. 1). This will also make it abundantly clear that the narrowband 60 Hz activity is a separate signal from the broadband induced activity. Whilst no statistically significant clusters were found for the enhancement of broadband activity by the flicker stimulus, compared to the static grating (p = 0.43 for the ‘most significant’ cluster), Suppl Fig 1E-F show that also when removing the immediate 60 Hz flicker effect, the flicker stimulus significantly enhanced high-frequency activity (when not correcting for multiple comparisons), albeit the effect is considerably weaker compared to when not removing the 60 Hz activity. Such effects have been described previously (Herrmann, 2001). Please note that the error-bars (in all figures shown here) reflect the dispersion across participants, whereas this is a within-subject design.

#### Flicker entrained narrowband 60 Hz activity

Finally, Figure 3I and J show the results of the comparison of the narrowband response to flicker and static stimuli. This comparison was based on averaged complex Fourier spectra, i.e. looks at the phase-locked (or ‘evoked’) response to stimuli. The thresholded topography (cluster-level p = 0.00025, the smallest possible for our permutations) in Fig 3I shows, to some surprise, clear maxima of signal enhancement: one in posterior occipital cortex, blending into more anterior parts of inferior occipito-temporal cortex and one in medial parietal cortex, just dorsal of the precuneus. The same topography was in fact also observed when averaging the absolute Fourier-coefficients, i.e. when looking at the induced narrowband response (i.e. not just the phaselocked component) -demonstrating a high level of consistency. Figure 3J shows the comparison of the spectra for flickered and unflickered stimuli in these significantly modulated cortical grid points (Fig 3I). Graphs of individual spectra, showing the total response (not just evoked, phaselocked activity) are provided in Fig 4B – showing that the measurement of the specific flicker-response does not depend on making use of the consistent phase of the flicker stimulus, but is a signal that can be measured in each individual participant with a high signal to noise ratio (SNR).

#### Individual profiles and frequency-shift following flicker ‘entrainment’

We next turned to the question how the enhancement of broadband induced gamma-band oscillations by the external 60 Hz flicker might depend on a participant’s intrinsic visually induced gamma-frequency (for the given stimulus). In Figure 4A and B we show the individual spectra of broadband and narrowband activity in the calcarine sulcus (Fig 3) for both conditions (flicker and static) for each corresponding participant. It is evident that the shape of the spectra (of the broadband induced gamma-response) differs considerably between participants, some showing several peaks, others one broadband peak and others show no induced gamma-peak at all (S5 and S9 in the static condition).

We next plot the peak frequencies (see methods section for details on how these were determined) in the two conditions against each other (Figure 4C). Each circle corresponds to one participant where the x-coordinate corresponds to their peak-frequency during the static condition and the y-coordinate corresponds to their peak-frequency during the flicker condition. In order to avoid overlays (due to the discrete nature of the frequency resolution), some random noise was added to both frequencies in this display. It is apparent that some participants shift their peak-frequency to the flicker-frequency (60 Hz) in the flicker condition (marked in red in Fig 4C), as one might expect, whereas others do not. Specifically, those that are located on the diagonal and are marked in blue retain their intrinsic frequency (or at least do not adopt the 60 Hz peak frequency) also during the flicker condition. Participants whose intrinsic frequency is 60 Hz are of course ambiguous in this respect and are marked in black. Note, however, that the intrinsic frequency (or its distance from 60 Hz) during the static condition does not appear to determine whether such a frequency shift occurs in that there are participants with intrinsic frequencies well below 60 Hz that either shift or do not shift their frequency or do not shift (no statistically significant difference in static frequency or distance from 60 Hz; all absolute(t) < 1, p > 0.39). This stands in contrast to what one might expect from the viewpoint of resonance in oscillatory systems.

#### Resonance phenomena

We then took this further and investigated whether the increase in power by the flicker frequency would depend on the proximity of the 60 Hz extrinsic flicker to an individual participant’s native gamma-peak frequency, as one would expect on the basis of resonance properties of oscillatory systems. Figure 4D shows this by plotting on the y-axis the peak-amplitude-enhancement of the broadband induced response by the flicker-stimulus against the difference of an individual’s peak frequency (measured during static condition) from the 60 Hz flicker frequency. This suggests the presence of the expected statistical relationship, that those participants with an intrinsic frequency near the flicker frequency would show a stronger enhancement of their broadband induced gamma-response by the flicker stimulus. The correlation analysis between the absolute distance of the intrinsic peak frequency from 60 Hz and the peak-power-difference between flickered and static condition showed that this was indeed the case for induced responses (r = −0.73, p < 0.01) and is shown in Figure 4D, but this relationship was not significant for the narrowband 60 Hz activity (r = −0.43, p = 0.12, not shown). There was no difference in either peak oscillatory amplitude for the two groups (‘shifters’ vs ‘non-shifters’, all absolute(t) < 1.1, p > 0.3).

#### Phaselocked entrainment

We next looked at the extent to which the flicker-stimulus would entrain both broadband gamma-activity, as well as the narrow-band 60 Hz activity, and whether this would depend on a) the distance of the native frequency from the flicker frequency and b) whether the induced gamma-peak-frequency shifted to the flicker frequency or not.

Supplementary Figure 2A shows the outcome of this analysis for the broadband induced gamma-activity (the frequency analysis was adapted for this purpose, see methods section), suggesting a high degree of entrainment even of the broadband-gamma oscillations that increases with time. It can, however, not be ruled out that this metric is influenced by the direct flicker effect (which is superimposed on the broadband oscillations, as we have shown). It is, to our knowledge, impossible to analytically distinguish between these two scenarios. The strength of induced gamma-oscillations are decreasing over time (Suppl Fig 2B), whereas the strength of the narrowband-flicker rhythm remains stationary (another ad-hoc analysis for the flicker induced activity with a temporal resolution of 1 s was run here). The increase of the broadband phaselocking value over time could therefore either be due to genuine increased phaselocking of the intrinsic gamma-rhythm to the flickering stimulus, or simply reflect the reduction of the impact of the broadband gamma-rhythm on the phase estimate (thereby enhancing the relative weight of the direct flicker response).

It was to be expected that the narrowband-flicker-related activity would show strong phaselocking to the stimulus timecourse, hence we do not show this in greater detail here. Figure 4E thus shows the phaselocking values for the narrowband 60 Hz frequency-analysis in the same way as Fig 4D, as a function of the spectral difference and for the subgroups and Figure 4F shows this for the 60 Hz frequency bin of the broadband frequency-analysis (grandaverage shown in Suppl Fig 2). Please do note, again, that these two estimation “points” have radically different spectral integration properties, the first one integrating over a width of approximately 0.25 Hz, the second one over a width of approximately 20 Hz – each of them with a real-world relation to different signals of different neurophysiological origin. Do note the remarkably high phase-locking values observed here (between 0.3 and 0.9). There was, however, neither a statistically significant correlation between the phaselocking-value for either rhythm and intrinsic frequency (all absolute(r) <= 0.4, p > 0.1), nor was there a significant difference between the two subsamples (all absolute(t) < 1, p > 0.5). This suggests that the questions of whether a) the frequency of the induced gamma-rhythm shifts to the flicker rhythm or b) the amplitude of the induced rhythm is strongly enhanced by the flicker is not strongly related to the degree by which (part of, namely the 60 Hz ‘component’) the native induced rhythm is phase-entrained by the stimulus.

#### Relevance of either rhythm for behaviour

Previous research has shown that the strength of visually induced gamma-oscillations predicts reaction time in similar change detection (M. Bauer et al., 2014; Hoogenboom et al., 2010; Womelsdorf et al., 2005) or even simple reaction time tasks (M. Bauer et al., 2009). A key question was therefore whether we could replicate these results in these data and which rhythm in visual cortex would be more correlated to behaviour – an indication of their functional significance for downstream processing.

In Figure 5 we show the results of the statistical comparison between fast and slow trials during the different experimental conditions and in different participants. Figure 5A shows the (thresholded for omnibus statistical significance) topography of the statistical comparison for static stimuli, correcting for multiple comparisons (p = 0.01). Figure 5B shows the spectrum of this comparison (uncorrected) in significantly modulated grid-points. These results are nearly identical to those published by (Hoogenboom et al., 2010), thus replicating these effects in this dataset on source level (we had not been able to replicate this using sensor level analysis on planar gradients before). Figure 5C shows the same contrast for the flicker condition – and evidently shows a clear absence of this effect in the static condition. The cluster-permutation analysis did not reveal any statistically significant cluster for the same comparison in the flickered condition. In an attempt to further distinguish between broadband induced and flicker entrained 60 Hz activity, we computed the same analyses on activity after removing the 60 Hz activity. The results are shown in Supplementary Figure 3. As a control, we also did this for the static condition and effectively replicated the analysis presented in Fig 5A-B, see Suppl. Fig 3A and B (p = 0.047). Like in the main analysis without the 60 Hz component removed, no cluster was observed for the flickered condition. The time-frequency plot of the statistic when using the grid points that revealed significant modulations in the static condition is shown in Suppl. Fig. 3C.

Figure 5D shows this now for the narrowband activity -only in the flicker condition (p = 0.02); there are, unsurprisingly, no effects in this metric in the static condition. This topography shows the same contrast obtained with the same method as in Fig 5A now for the narrowband 60 Hz activity (flicker only). Note the similarity of this narrowband 60 Hz topography in the flicker condition with the broadband induced topography in the static condition (Fig 5A).

Figure 5E shows the power spectra for fast and slow trials (again, only in the flicker condition; there is no difference in the static condition for this activity), clearly indicating that, in the flicker condition, it is the narrowband 60 Hz response that is ‘correlated’ with behaviour. This is thus a separate response that is mostly invisible in the broadband response (Fig 5C).

This may initially appear counter-intuitive for those familiar with frequency analyses, in that one might, at first sight, expect such a response to show up also in Fig 5C. However, as explained in some detail in the methods section and above, this (inherently!) narrowband neuronal flicker response easily gets overshadowed in the MEG sensors by more broadband background activity -if not specifically adjusting the frequency resolution to measure this particular activity. This is not only true for the very wide spectral smoothing with multi-taper analyses, but applies generally, as witnessed in our very own experience previously, e.g. (Kamphuisen et al., 2008). There is thus no contradiction between Figure 5C and E, these represent genuinely different brain activities (measured with different weightings in each case). Finally, the above result (contrast fast vs slow trials) on narrowband activity in the flicker condition is also found if not only looking at the phase-locked part but the total narrowband induced response, albeit the contrast is then weaker, presumably due to the reduced SNR.

Finally, we investigated whether there might be any inter-individual differences in the relation between the broadband induced vs narrowband evoked rhythms and behaviour. To this end, we investigated those participants shifting their intrinsic peak-frequency to the flicker show a different pattern compared to those who do not shift their peak-frequency. Figure 5F shows this for the broadband induced activity in the flicker condition. Those participants that shift their peak frequency to the flicker (red in Figures 4C-F, as well as red line here), i.e. those participants where the entrained flicker rhythm appears to gain larger control of the intrinsic gamma-rhythm, show tentatively weaker induced gamma-responses during fast compared to slow trials during the flicker condition, i.e. they show an inversion of the pattern seen in Fig 5A,B. By contrast, in those participants who retain their peak frequency also during the flicker stimulus (blue circles in Fig 4 and blue line here), the relationship between broadband induced gamma-oscillations and RT remains existent. This difference between the two sub-samples is highly significant (black line in Fig 5F, p<0.01 around 80 Hz). No clear difference between the two subsamples was found on the narrowband response – both showed highly significant differences as a function of RT during the flicker condition there.

To summarize this last section, across the sample broadband induced gamma-band activity in lateral occipital cortex correlates with RT for unflickered stimuli, suggesting the relevance of this signal for overt decision making. However, when a grating is flickered, induced broadband gamma-activity remains intact and is in fact enhanced, but, across the sample shows no relationship with RT; instead the narrowband flicker-entrained activity becomes dominant with respect to its relevance over behavioural control. In those participants though, where the induced gamma-band activity appears to be less controlled by the entrained rhythm, both rhythms simultaneously are correlated with behaviour.

## Discussion

We show here the co-existence of two largely separate rhythms during the presentation of 60 Hz flickered visual gratings: a rather narrowband 60 Hz rhythm, measured in each participant and the classical broadband visually induced gamma-rhythm that is also present during ‘normal’, i.e. unflickered presentation of visual gratings. Whilst all participants showed some extent of phase-entrainment around 60 Hz and we observed a general enhancement of broadband oscillations in the flicker condition, in several participants the dominant frequency of the intrinsic visually induced gamma-oscillation remained unaltered and the broadband nature (approx. 20 Hz) of this oscillation generally remained intact. This, besides other facts, indicates the co-existence of two rhythms, rather than the control or mere amplification of the intrinsic rhythm by an external force. Furthermore, the results suggest that these two rhythms may contribute independently to behaviour, potentially even in some competitive fashion, with strong and systematic group differences regarding the relevance of either rhythm for the propagation of sensory information to higher level processing stages. Since this type of paradigm has frequently been used and is being used to probe the causal role of brain oscillations for neural information processing and cognition, the results raise important questions about the validity of this approach and the interpretation of the results of such studies.

The flickered stimulus proved to very robustly entrain a 60Hz rhythm in visual cortex in each individual participant in our study, even if not just looking at the phaselocked steady-state-evoked-response, but the total power. This finding is not entirely trivial given human EEG studies (Herrmann, 2001) that reported a sharp decline of the amplitude of entrained flicker rhythm with frequency, in particular for frequencies above 30Hz. A key factor for the sensitivity of an analysis to detect such rhythms is the spectral resolution. Not only at such high frequencies, but already at much lower frequencies, the risk is high that the very narrowband activity that is entrained by flicker stimuli (Srinivasan et al., 1999) is otherwise overshadowed by unspecific ongoing physiological activity and environmental noise (an experience collected over the years and in particular during this study (Kamphuisen et al., 2008)). Importantly, the presence of the two rhythms is not just an artefact of the two frequency analyses run here either, as a naïve reader might come to think. As shown, particularly in Figure 3, the flicker entrained rhythm genuinely has a very narrowband response, always at 60 Hz, whereas the individual intrinsic gamma-rhythms have different frequencies. The broadband nature of the induced rhythm is very well known from hundreds of studies and the spectral features known to depend on both individual (in particular also genetic) features (Muthukumaraswamy, Edden, Jones, Swettenham, & Singh, 2009; Pinotsis et al., 2013; van Pelt et al., 2012) as well as stimulus properties (Hermes, Miller, Wandell, & Winawer, 2015; Orekhova et al., 2015). Our stimulation setup only allowed to stimulate at a fixed frequency of 60 Hz, which is known to be approximately the average gamma frequency in humans (Hoogenboom et al., 2010) and was also the average frequency in this study, when taking the initial gamma-band response into account (and somewhat lower when focusing on the stationary part of extended grating exposition).

Importantly, besides the mere co-existence of the two rhythms, we did find that the stimulus-entrained (narrowband) rhythm on average across the whole sample appeared to potentially have partially replaced the ‘causal relevance’ (we will discuss this term and this interpretation that may at first appear as a bit of a stretch below) of the induced gamma-rhythm for sensori-motor processing: whereas for non-flickered stimuli, faster reaction times were accompanied by stronger induced gamma oscillations, no such difference was observable (across the group, see next paragraph for a discussion of inter-individual differences) in the flickered condition. Instead, for flickered stimuli, faster reaction times were accompanied by enhanced 60 Hz narrowband entrained oscillations. Whilst the observation of such a mere correlate of course precludes any inferences on causality, since many other signatures of brain activity are likely to contribute to faster reaction times – this double dissociation is of particular interest that makes an important difference in that respect. Whatever the nature of the mechanistic process that drives this correlation of enhanced signal amplitude with reaction time, the effective replacement of one signal correlate with another one in different experimental conditions (in the absence of the diminishing of the broadband induced rhythm) can, in our opinion, not reasonably be explained by anything but the fact that these signals are playing instrumental roles in the efficient propagation of visual information to motor preparation stages: the entrained rhythm during the flickered stimulus and the induced rhythm during non-flickered presentation.

Participants were not visually aware of the 60 Hz flicker, hence it is highly unlikely that they were attending selectively to the narrowband 60 Hz signal (besides that such a process is unknown to us). If, for instance, variations of attention or vigilance during the task may jointly lead to faster reaction times (under enhanced attentive state) and enhanced broadband and narrowband oscillatory amplitudes (for all of which there is ample evidence, see (M. Bauer et al., 2014; Muller et al., 1998; Posner, Snyder, & Davidson, 1980)), which is an entirely plausible scenario (that neither speaks against nor strictly in favour of a causal role of these oscillations), one would expect to see enhanced signals in induced and entrained activity during the flicker condition. However, this is not what we observe here. We can only plausibly interpret this double dissociation in such a way that the induced gamma-oscillations that appear to have an important role in signal propagation during the unflickered grating lose out against the signal of the entrained rhythm that appears to have higher significance for efficient sensorimotor processing during the flicker condition. As we discuss further below, the most parsimonious explanation for the results of this and other studies may be that entrained and induced rhythms are generated in (at least partially) different neuronal populations with each having different importance for information processing during these conditions.

Further in this respect, we show that there are considerable inter-individual differences in the entrainment of gamma-oscillations and that this correlates with the significance of the intrinsic rhythm for sensorimotor propagation or behavioural control. In some participants, the intrinsic gamma-oscillations were under “tighter control” by the external force than in others, in that their intrinsic gamma-rhythm shifted to the peak-frequency, whereas others did not show this behaviour. Crucially, this appears to be largely independent from the match of the frequency of the external stimulus to the intrinsic gamma-peak frequency. Whilst we describe in the paragraph above that across our entire sample, in the flicker condition the induced rhythm was not enhanced for faster RT, this was only true for those particpants whose intrinsic gamma-frequency shifted to the flicker frequency (Fig. 5F). In those participants, in fact we observed a reduced gamma-amplitude with fast RT in the flicker condition, whereas in those that did not shift, the broadband induced gamma-oscillations remained (somewhat) enhanced for faster RT.

We do not know what may cause these interindividual differences – please note that the two subsamples cannot be differentiated in terms of their overall signal amplitudes, not even in the strength of the entrainment of either rhythms by the flicker stimulus (see Figure 4 and statistical analyses in the results section). However, and most importantly, they may provide an explanation for the highly inconsistent findings concerning the use of rhythmic brain stimulation – by either sensory stimuli (Dakin & Bex, 2002; Elliott & Muller, 2000; Fahle & Koch, 1995; Kiper et al., 1996; Leonards, Singer, & Fahle, 1996) or direct brain stimulation (Heroux, Loo, Taylor, & Gandevia, 2017; Kim et al., 2014; Veniero, Benwell, Ahrens, & Thut, 2017).

We found that the degree by which the induced visual gamma-oscillations were enhanced by the 60 Hz flicker, depended on the match of the flicker rhythm to the intrinsic gamma-band activity (in response to the non-flickered stimulus). Such resonance effects are to be expected on the basis of the temporal dynamics of most oscillatory systems (Pikovsky et al., 2001). However, another, related study (Duecker, Gutteling, Herrmann, & Jensen, 2020), that overall confirmed the more general conclusions of this paper (of a co-existence of directly entrained flicker response with the induced gamma-oscillations), found no such resonance effects and also claim that their flicker stimuli did not cause phase entrainment.

However, besides the fact that their figures 2 and 3 suggest that there may in fact be some resonance effect (overshadowed by a 1/f dropoff), their study differed from ours in several important ways: They superimposed the flicker (sinusoidal amplitude modulation of luminance) over a circular moving grating as one stimulus and over a circular patch of luminance as another stimulus. Regarding the first stimulus, this causes two types of dynamic modulations that (smaller) receptive fields in early visual cortex will experience: the spatial displacement (movement) of the grating over time and the amplitude modulation of the luminance values, leading to a mixture of different frequencies, and reduced efficacy of the flicker since a given receptive field will only temporarily experience the flicker (while the ‘white stripes’ are in its receptive field). Regarding their patch stimulus, this will trigger neurons with very different spatial (and potentially) temporal frequency preferences and such stimuli are generally thought to create weaker responses than for instance checkerboard or grating stimuli.

At the same time, they had access to a visual projector offering a very high frequency (vertical refresh of 1.44 kHz) display that allowed them to create truly sinusoidally modulated ‘photic drive’ signals at several frequencies. It is of course also possible, that neurons in visual cortex do not respond as strongly to smooth changes in amplitude at such high frequencies, compared to our alternating frames design (please note that we still got a very narrowband 60 Hz oscillatory response from this). We consider it thus likely that their differing stimuli may be responsible for the relative lack of resonance seen, compared to our study.

They further claim that there was no phase-entrainment of the induced gamma-rhythm by their photic drive stimulus. This stands, again, in contrast to what we find. We find, very strong phase entrainment effects, not only (as more trivially to be expected) in the narrowband 60 Hz response directly imposed by the flicker. We also find rather high phaselocking values to the flicker in the broadband induced gamma at 60 Hz. We cannot with any certainty distinguish whether this simply picks up the directly entrained flicker signal, but we consider this to be less likely for two reasons: 1) the direct flicker entrained activity does not influence the power estimates from our multitaper method with its broad spectral integration properties much (see for instance Figures 4 and 5 for radical dissociations between these two measures, where a strong 60 Hz component in the narrowband analysis is totally invisible in the multitaper estimates); 2) the phaselocking-value of the broadband induced gamma increases over time, whereas the direct 60 Hz flicker response amplitude remains constant. Ultimately, this latter question might only be answered with methods that distinguish between individual neurons. Generally, our two studies come to overall similar conclusions: that there is a coexistence between the two rhythms with only relatively weak interactions, if any. Our study is going further in that we provide strong evidence that in the presence of the flicker rhythm, this activity seems to become dominant with respect to signal propagation to higher level decision stages and may in fact ‘replace’ the importance of the intrinsically generated gamma-rhythm.

Interestingly, our previous TMS study entraining beta-oscillations over motor cortex (Romei et al., 2016) has provided results that tie in with these questions regarding resonance and phase-entrainment: The amplitude of the effect of an extrinsic rhythm on brain activity depends on the match to the intrinsic frequency. However, in this study we found no lasting effect of phase-entrainment on the cortical beta-oscillation, as one would expect if the intrinsic rhythm had been entrained. A likely explanation for this is that ‘entrainment’ of brain activity through rhythmic stimulation (sensory or TMS) may affect at least partly different neurons than those involved in generating the intrinsic rhythm. The resonance effect (in terms of amplitude enhancement as a function of frequency match) might then be explained by these neurons having similar spectro-temporal filtering characteristics.

The results presented here are of course to some extent specific for visual flicker stimulation at gamma-frequencies. Entrainment by flicker stimulation has in the past, at least for alpha-oscillations, been shown to be a highly potent way of entraining brain oscillations with “long”-lasting phase-entrainment (Spaak, de Lange, & Jensen, 2014), whereas our previous study of entraining brain oscillations through direct brain oscillations using TMS caused no such long-lasting phase-entrainment (Romei et al., 2016): the phase of the beta-oscillation after the end of TMS stimulation showed no relationship to the TMS train.

Taken together these results suggest that studying the causal role of gamma-oscillation using external stimulation paradigms remains a difficult issue. Ours and the study by (Duecker et al., 2020) show that entrainment of brain activity, even when the driving frequency is matched to the individual intrinsic frequency, does not necessarily manipulate the intrinsic rhythm strongly and the effect of such manipulation on neural information processing may be such that it might bypass or even attenuate the intrinsic rhythm. Here we have shown that there is considerable heterogeneity between participants that may partly explain the lack of consistency in studies aiming to study the causal role of brain oscillations through external entrainment. It will be of interest to investigate this for tACS and other brain stimulation studies, although the confound of applied electric and natural rhythms makes these analyses naturally more complex.

## Acknowledgements

The experiment was conceived and the data were recorded in 2014 with the help of another undergraduate student, Xinyan Zhou. Markus Bauer received funding from a University of Nottingham Research Fellowship.

