## Supplementary Figure for "Can the causal role of brain oscillations be studied through rhythmic brain stimulation?"

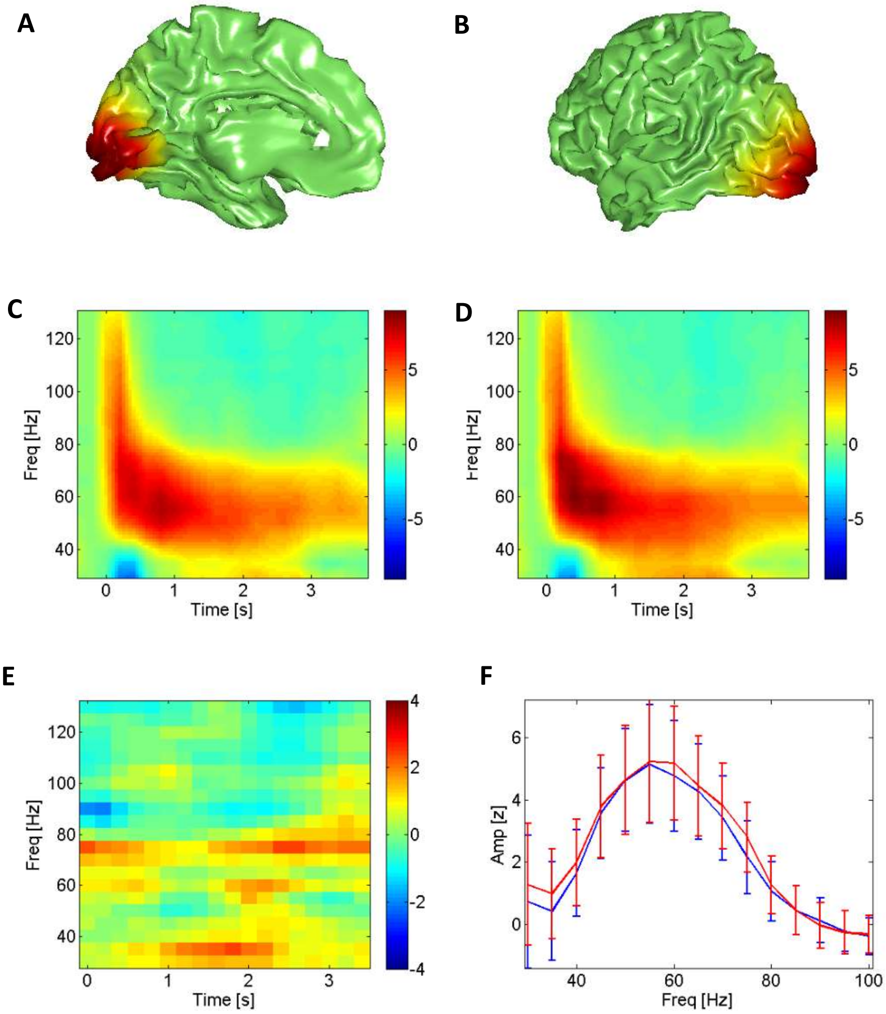


**Supplementary Figure 1: Induced gamma oscillations after removal of narrowband 60 Hz activity**
This figure shows the results of the broadband (multitaper) frequency analysis after removal of the narrowband 60 Hz activity (bandstop filter 59.75-60.25 Hz). **A), B)** Topography of statistically significantly modulated grid-points (in static condition), in analogy to Fig 3 A,B (medial and lateral views of left hemisphere, respectively). **C)** TFR of unflickered condition after 60 Hz removal – note that there is no 'dip' in the PSD at 60 Hz due to the 20 Hz smoothing properties of the multitaper frequency smoothing. Colourbars represent t-values. **D)** Flicker induced gamma-oscillations after 60 Hz activity removed. **E)** Comparison of power-spectra in flicker vs static (paired t-tests) conditions in same grid points as in Fig 3E,F. The cluster-permutation-test, correcting for multiple comparisons did not yield (omnibus) significant differences. **F)** Overlay of the power spectral densities in analogy to Fig 3H

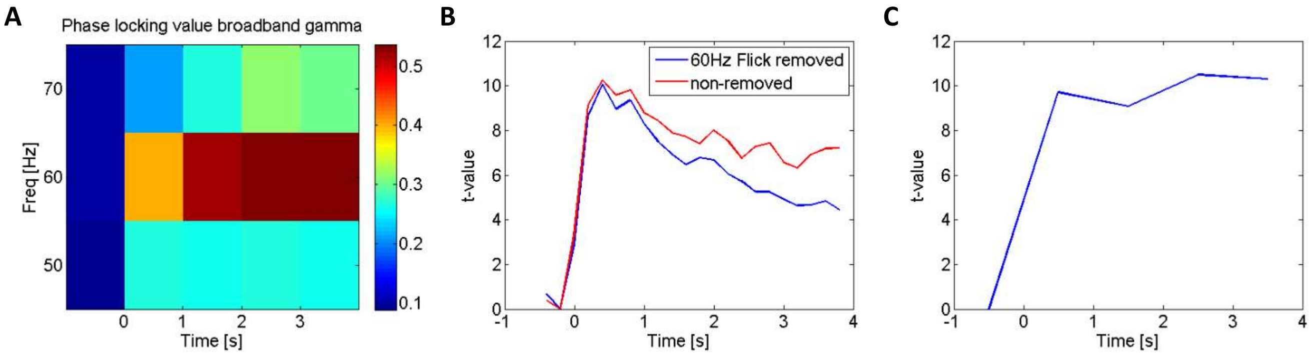


**Supplementary Figure 2: Broadband induced gamma phaselocking-values, in comparison to timecourse of broadband and narrowband induced power**

**A)** TFR of phaselocking value (PLV) of broadband induced gamma-oscillations to the photodiode signal in Flicker condition in calcarine sulcus (grandaverage of 'raw' PLV values). **B)** Timecourse of induced gamma-band activity at 60 Hz with 60 Hz activity removed (blue line) and without 60 Hz activity removed (red). **C)** Timecourse of narrowband 60 Hz activity (1 Hz bandwidth, 1 s time-windows centered on -0.5, 0.5, 1.5, 2.5 and 3.5 s post-stimulus) that was additionally run for this comparison only.

Gamma-band phaselocking over the course of the stimulation appears to increase, in contrast to the amplitudes of narrowband 60 Hz power which remains stationary. However, this might be partially due to decreased interference of the non-phaselocked broadband induced gamma, the amplitude of which decreases over time.

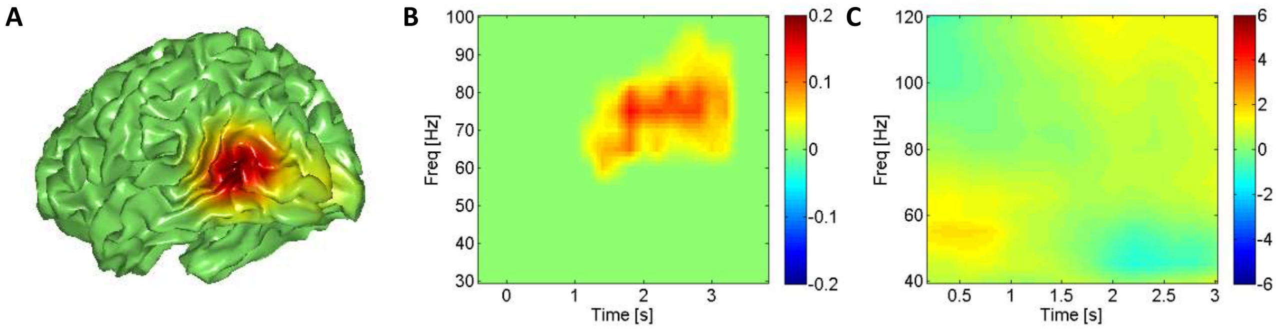


**Supplementary Figure 3: Fast vs slow trial induced responses in flicker and static condition, after removal of narrowband 60 Hz activity**

**A)** Topography of RT effect (thresholded for multiple comparisons) in static condition, in analogy to Fig 5A, but here after removal of 60 Hz activity. **B)** TFR of statistical comparison between fast and slow trials in static condition, after removal of 60 Hz activity (thresholded for multiple comparisons). Colourbar represents arbitrary units. **C)** Unthresholded TFR of comparison between fast and slow trials in flickered condition (60 Hz removed, otherwise analogous to Figure 5C), in the same grid points as in A). Colourbar here represents real t-values, no statistically significant effects were found. These analyses confirm the results of Figure 5.
